# Epidermal stem cell-derived extracellular vesicles induce fibroblasts mesenchymal-epidermal transition to alleviate hypertrophic scar via the miR-200s/ZEBs axis

**DOI:** 10.1101/2025.01.27.635177

**Authors:** Miao Zhen, Juntao Xie, Rui Yang, Lijuan Liu, Hengdeng Liu, Xuefeng He, Suyue Gao, Junyou Zhu, Jingting Li, Bin Shu, Peng Wang

## Abstract

Hypertrophic scar (HS) is a prevalent yet unresolved wound healing complication characterized by persistent hyperactive and proliferative fibroblasts, leading to excessive extracellular matrix (ECM) synthesis and collagen contraction. Our previous studies have identified epidermal stem cells (ESCs) as critical for wound healing and HS remodeling, with its extracellular vesicles (EVs) playing a vital role. However, the specific mechanisms remain unclear. In this study, we first discovered that ESC-EVs could effectively induce the mesenchymal-epidermal transition (MET) of HS fibroblasts (HSFs) and inhibit their biological activity. Furthermore, by next-generation sequencing and multiplexed CRISPR/Cas9 system, we elucidated that this therapeutic effect is mediated by the miR-200 family (miR-200s) encapsulated in ESC-EVs, which targeted and inhibited ZEB1 and ZEB2 in HSFs. This vital role and mechanism have been thoroughly validated in both in vitro cell experiments and in vivo rat tail HS (RHS) models. These findings not only shed light on a previously unidentified mechanism of ESC-EVs for HS, but also provide potential novel targets and strategies for its precise treatment.

## 1 INTRODUCTION

Hypertrophic scar (HS) emerges as a frequent consequence following dermal insults such as burns, trauma, puncture wounds, acne, surgical incisions, and vaccinations (Ogawa, 2017). Clinically, HS presents patients with tremendous challenges by various symptoms, including pruritus, pain, paresthesia, erosion of underlying bone structure, and restriction of joint mobility (Berman *et al*., 2017). However, despite the array of therapeutic approaches available, from corticosteroids and pressure therapy to laser treatments and surgical excision, the therapeutic effect of HS remains unsatisfactory (Rabello *et al*., 2014). Urgent efforts are warranted to identify precise intervention targets and develop novel strategies for HS.

Recent studies have demonstrated that epithelial-mesenchymal transition (EMT) is crucial in tissue fibrosis and scar formation (Youssef and Nieto, 2024). EMT represents a ubiquitous biological process where polarized epithelial cells undergo loss of cell polarity and intercellular adhesion and acquire a mesenchymal cell phenotype, which culminates in heightened deposition of extracellular matrix (ECM), augmented resistance to apoptosis, and intensified migratory and invasive potential (Boyer *et al*., 2020). Although EMT is necessary for proper re-epithelialization in routine wound healing, the uncontrolled continued transition from epithelial cells to hypertrophic scar fibroblasts (HSFs) eventually leads to aberrant ECM deposition, collagen contraction, and disrupted structural integrity (Machesney *et al*., 1998). Therefore, inhibiting epithelial cells’ EMT or even inducing HSFs mesenchymal-epithelial transition (MET) should be highly efficient in alleviating HS.

Emerging evidence suggests that extracellular vesicles (EVs) regulate EMT or MET and mediate the development of various fibrosis diseases (Jiang *et al*., 2022). As a crucial subset of epithelial cells and progenitors of diverse epidermal cell lineages, Epidermal stem cells (ESCs) possess the capacity to proliferate and differentiate into various epidermal cell types, usually showing impeccable efficiency for wound healing and the mitigation of scar formation than other kind of stem cells (Blanpain and - Fuchs, 2006; Moore and - Lemischka, 2006). Encompassing a diverse array of proteins, DNA, and RNA that mirror the physiological profile of ESCs, EVs derived from ESC (ESC-EVs) mimic the functional properties of ESCs and present considerable potential as a substitute for cellular therapy. Encouragingly, our recent studies have discovered that ESC-EVs have an ESCs’ identical therapeutic effect in wound healing and scar prevention (Wang *et al*., 2017a, 2019, 2022a), and exhibited a potential to modulate EMT and MET. However, its crucial mechanism urgently needs to be further explored.

In this study, we established an efficient ESC-EVs isolation and application system and thoroughly validated their effects on MET induction and biological function inhibition of HSFs in vitro and in vivo. Furthermore, using next-generation sequencing and multiplexed CRISPR/Cas9 system, we delineated that ESC-EVs induced HSFs’ MET mainly through miR-200s/ZEBs axis. This innovative mechanistic insight suggests a promising avenue for future therapeutic strategies targeting HS and other skin fibrosis diseases.

## 2 MATERIALS AND METHODS

### 2.1 Cell isolation and culture

Human skin tissue samples were obtained from The First Affiliated Hospital of Sun Yat-Sen University, and ethical approval and informed consent were obtained. ESCs and fibroblasts (FBs) were isolated from human foreskins, while hypertrophic scar fibroblasts (HSFs) were isolated from human hypertrophic scars as described (Wang *et al*., 2022b). Briefly, subcutaneous tissues and fat were removed before the skin was disinfected and incubated overnight with DispaseL (3mg/ml; D4693-1G, Sigma-Aldrich) in keratinocyte serum-free medium (Epilife, MEPI500CA, Gibco) with EDGS (S0125, Gibco) at 4L to separate the epidermis from the dermis. After that, the epidermis separated from the foreskins was digested with prewarmed Trypsin/EDTA (0.25%, Gibco) at 37L for 10 minutes to obtain ESCs, which were resuspended in Epilife and seeded at a density of 10^5^ cells/cm^2^ in 100 μg/ml type L collagen-coated culture dishes. Cells adhering within 10 mins were selected according to the rapid attachment method(Kim *et al*., 2004). Similarly, the dermis was digested with Collagenase IV (3mg/ml, C6745-1ML, Corning) and Hyaluronidase (2mg/ml, 1141GR001, Biofroxx) in Dulbecco’s modified Eagle medium (DMEM, C11995500BT, Gibco) at 37L for 2 hours to obtain FBs and HSFs, which were cultured in DMEM supplemented 10% fetal bovine serum with 5% CO_2_ at 37L. All cells were cultured in cell factories to obtain sufficient amounts of EVs. Cells at the 3-5 passages were used for this study.

### 2.2 EVs isolation, characterization, and treatment

EVs were isolated from the culture medium of cells using a differential ultracentrifugation method[(Lin *et al*., 2020)]. For the collection of conditioned medium, 80% confluent ESCs or FBs were washed with PBS before being cultured with serum-free media for 2 days. Briefly, the medium was centrifuged at 300×g for 10 min and 2000×g for 10 min at 4L, followed by ultracentrifugation at 100000×g for 70 min at 4L on an Optima XPN-100 ultracentrifugation with an SW32Ti rotor (Beckman Coulter, USA). The isolated EVs were washed with PBS and subjected to secondary ultracentrifugation at 100000×g for 70 min at 4L. EVs were stored in PBS at −80L. The particle size distribution and concentration were measured with ZetaView (Particle Metrix, Germany), and the morphology was examined by electron microscopy (HITACHI HT-7700, Japan). The EV protein was quantified using a BCA protein assay kit (Thermo Fisher Scientific, USA). Western blotting was performed to detect positive markers Alix (1:1000, RGAB100-50, Rengenbio), CD9 (1:1000, ab92726, Abcam), CD81 (1:1000, RGAB105-50, Rengenbio), CD63 (1:1000, ab134045, Abcam), TSG101 (1:1000, ab125011, Abcam) and negative markers GAPDH (1:50000, 60004-1-lg, Proteintech) of EVs.

### 2.3 Cellular uptake assay of EVs

To evaluate EVs delivery, PKH67 membrane dye (MINI67, Sigma-Aldrich) was added to PBS at a concentration of 1μM and incubated with EVs (10μg) for 20min. The incubation was stopped with 0.5%BSA (A3311, Sigma-Aldrich) before washing repeatedly. The labeled EVs were then resuspended (10μg/ml) and incubated with HSFs for 24h. After that, the cells were fixed with 4% paraformaldehyde for Actin (2 drops/ml, R37112, Thermo Fisher) and DAPI staining and observed using a confocal laser scanning microscope (OLYMPUS FV3000, Japan).

### 2.4 Flow cytometric analysis (FCA)

To identify epidermal stem cells, about 10^6^ cells were collected, prepared as single-cell suspensions, and incubated with 2.5μg Fc block in 100μL PBS. Surface marker anti-CD49f-FITC (1:100, 313605, Biolegend, USA) was used to stain cells at room temperature for 30min. While for intracellular staining, cells were permeabilized and fixed with the intracellular staining kit (554714, BD Biosciences) and then washed and stained with antibodies anti-cytokeratin 15 (1:100, ab52816, Abcam) at room temperature for 30min. After washing, cells were incubated with second antibodies (1:500, 4408S, CST) and washed twice before resuspending. The stained cells were analyzed using a CytoFlex flow cytometer (Beckman, USA). Data were processed by FlowJo Software v10.8.1.

### 2.5 Cell viability assays

Cells were plated onto 24-wells at a density of 10^5^ cells/cm^2^ in a 37L and 5% CO_2_ incubator. After 24h, living and dead cells were detected using the Calcein/PI Cell Viability/Cytotoxicity Assay Kit (C2015S, Beyotime, China) in accordance with the manufacturer’s instructions. Then the cells were observed under an inverted fluorescence microscope (OLYMPUS IX83, Japan).

### 2.6 Collagen gel contraction assay

According to previous articles (Ngo *et al*., 2006; Huang *et al*., 2022), HSFs were resuspended in 1.8mg/ml collagen solution at a concentration of 2 × 10^5^/ml. The mixture of cellular collagen was seeded in 24-well plates (0.5ml/well) and incubated at 37L for 1h to coagulate. Then, 500μL of the conditioned medium was gently added to each well, and the gel was separated from the plates by gently running the sterile pipette tip along the gel edges.

### 2.7 Multiplex CRISPR/Cas9 system and lentivirus transfection

As previously described, we assembled customized lentiviral vectors expressing sgRNAs targeting hsa-miR-200b-5p, hsa-miR-429, hsa-miR-200c-5p, and hsa-miR-141-3p into a lentiviral vector (CV279, Genechem, China) that ultimately expressed active Cas9 by Golden Gate cloning(Kabadi *et al*., 2014; Yu *et al*., 2022). The epidermal stem cells were infected with lentivirus expressing multiplex CRISPR system. Stable clones were selected after 10 days using 1μg/ml puromycin.

### 2.8 Western blot analysis

Protein samples were lysate using RIPA lysis buffer supplemented with protease inhibitors (Calbiochem, USA). Equal amounts of proteins were loaded and separated by 10% SDS-PAGE and then transferred to polyvinylidene difluoride membranes (PVDF, Millipore, USA). The membrane was blocked with 5% BSA and then incubated with one of the primary antibodies: anti-alpha smooth muscle (1:1000, GB111364, Servicebio), anti-COL1A1 (1:1000, E3E1X, Cell Signaling Technology), anti-N Cadherin (1:1000, A19083, Abclonal), anti-cytokeratin 1 (1:500, PA5-113746, Invitrogen), anti-cytokeratin 15 (1:10000, ab52816, Abcam), anti-E Cadherin (1:500, ab1416, Abcam), anti-ZEB1 (1:500, ab181451, Abcam) and anti-ZEB2 (1:1000, AF5278, Affinity Biosciences) overnight at 4L. Afterward, the PVDF membranes were incubated with HRP-conjugated secondary antibodies at room temperature for 1h. The immunoreactive bands were visualized with the enhanced chemiluminescence kit (Sigma-Aldrich) and analyzed by densitometry using ImageJ software.

### 2.9 Fluorescence in situ hybridization (FISH)

The paraffin sections were prepared and fixed as previously described (BB, 2013). SweAMI probes against miR-200s were used (Servicebio, China). Staining was performed as written in the manufacturer’s protocol. Briefly, slides were treated with proteinase K (20μg/ml, G1234, Servicebio) for 20min and pre-hybridization for 1h at 37L before hybridizing with probes overnight at 37L. Then, the slides were counterstained with DAPI and scanned afterward.

### 2.10 RNA extraction and real-time PCR assay

Total RNA from cells was extracted using Trizol reagent (Thermo Fisher) and reverse-transcribed into cDNA using the PrimeScript RT reagent kit (Takara) according to the manufacturer’s protocol. Quantitative real-time PCR (qRT-PCR) reaction was performed with the Roche 480 system (Roche) using the LightCycler 480 SYBR Green I Master Mix (Roche). In contrast, miRNA was extracted using Trizol reagent (Thermo Fisher), reverse-transcribed, and performed qRT-PCR using All-in-One miRNA qRT-PCR Detection Kit (QP115, GeneCopoeia). The relative mRNA expression levels of target genes were calculated using the 2^−ΔΔCt^ method.

### 2.11 *In vitro* immunofluorescent staining

A total of 10^4^ HSFs were resuspended in 500μl DMEM with 10μg/ml ESC-EVs, FB-EVs, or PBS and added into one well of an eight-well removable glass chamber slide (Nunc Lab-TekLChamber Slide System, Thermo Fisher Scientific). After 24h, cells were fixed with 4% formaldehyde, permeated with 0.1% Triton X-100 (Solarbio, China) for 15 min, and blocked with 2% BSA for 1h. Next, the cells were stained with primary antibodies, including anti-alpha smooth muscle (1:500, GB111364, Servicebio), anti-COL1A1 (1:100, E3E1X, Cell Signaling Technology), anti-N Cadherin (1:200, ab98952, Abcam), anti-cytokeratin 1 (1:100, PA5-113746, Invitrogen), anti-cytokeratin 15 (1:50, ab52816, Abcam), anti-E Cadherin (1:50, ab1416, Abcam), anti-ZEB1 (1:200, ab181451, Abcam) and anti-ZEB2 (1:100, AF5278, Affinity Biosciences) overnight at 4L. Cells were washed and incubated with secondary fluorescence antibodies (1:2000, 4408S, 4409S, 4412S, 4413S, Cell signaling Technology) for 1h and DAPI for 10 min at room temperature. Then the cells were observed under an inverted fluorescence microscope (OLYMPUS IX83, Japan).

### 2.12 miRNA sequencing analysis of EVs

After total RNA was extracted by TRIzol, the RNA molecules in a size range of 18–30nt were enriched by polyacrylamide gel electrophoresis (PAGE). The transcripts per million reads were used for the calculation of gene expression, and genes with |log2 (fold change)| ≥ 1 and *p*<0.05 were considered statistically significant. Target genes of each miRNA in EVs were predicted using publicly available algorithms multiMiR (Ru *et al*., 2014). Kyoto Encyclopedia of Genes and Genomes analyses and Gene Ontology analyses were performed to elucidate the functions and enriched pathways of statistically significant genes.

### 2.13 *In vivo* model of hypertrophic scars

Male Sprague–Dawley rats aged 8 weeks were purchased from the Ruige Biological Technology Corporation (Guangzhou, China). All experimental procedures were approved by the Ethics Committee Board of the First Affiliated Hospital of Sun Yat-Sen University and performed in accordance with the NIH Guide for the Care and Use of Laboratory Animals. Briefly, a 6×6 mm full-thickness skin defect was created on the dorsal side of the rat tail. Then the wound site was stretched by attaching a 3 cm stainless steel ring to the ventral side of the tail. After 3 weeks, RHS had been successfully epidermalized. Thereafter, 12 rats were randomly assigned into three groups: the ESC-EVs group (200μl, 10μg), the FB-EVs group (200μl, 10μg) and the control group (PBS, 200μl). We administered weekly subcutaneous injections to the scar. On day 28, all rats were sacrificed and the scar tissues were surgically removed for experiments (Zhou *et al*., 2019).

### 2.14 Histological examination

Skin tissue was fixed overnight in 4% paraformaldehyde at room temperature and embedded in paraffin. Afterward, 5-μm-thick slices were cut out. Some of them were stained with hematoxylin-eosin (H&E) and Masson and analyzed using a microscope (Kfbio, China). Others were blocked with goat serum and then stained with primary antibodies, including anti-alpha smooth muscle, anti-COL1A2 (1:200, A5786, Abclonal), anti-N Cadherin, anti-cytokeratin 14 (1:200, A15069, Abclonal), anti-E Cadherin, anti-ZEB1 (1:200, ab181451, Abcam) and anti-ZEB2 overnight at 4L. Sections were washed and incubated with HRP-conjugated secondary antibodies (1:2000, ab6728, ab6721, Abcam) for 1h, fluorochrome (15X, ASOP520, ASOP570, Asbio) for 10min, and DAPI for 10min at room temperature. Slides were scanned afterward (Pannoramic MIDI, 3DHISTECH, Hungary) and analyzed using CaseViewer 2.4.

### 2.15 Statistical analysis

All data are presented as the mean ± SD. Comparisons of changes in levels of signaling component expression between control and experimental groups at the same time point were conducted by using the Student’s t-test. The differences between the groups at different time points were compared by one-way analysis of variance followed by the Bonferroni test. Statistical analyses were performed with GraphPad Prism 8.3.0 software (GraphPad Software, USA); *p*<0.05 was considered statistically significant.

**Table 1.**
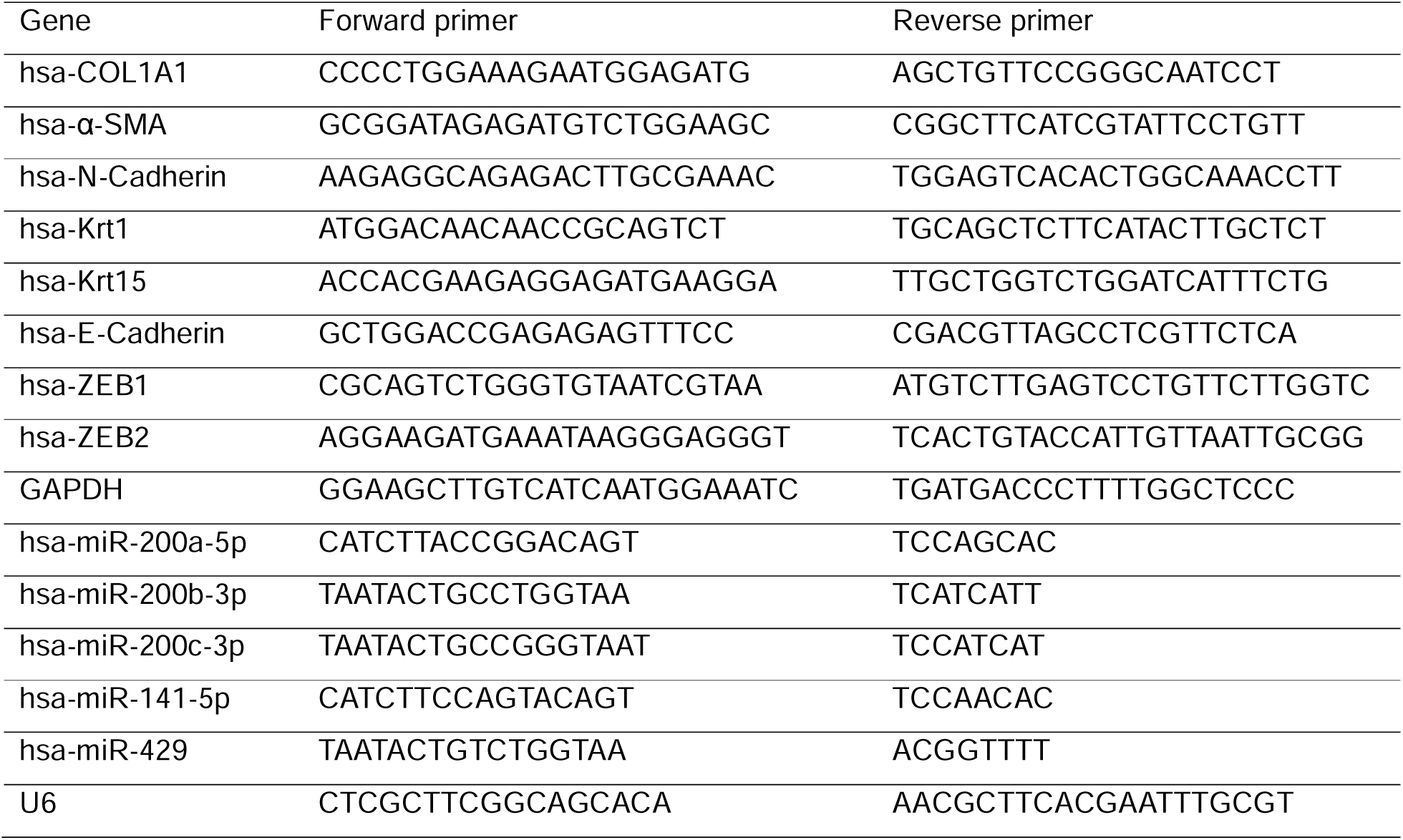
qRT-PCR primers

**Table 2.**
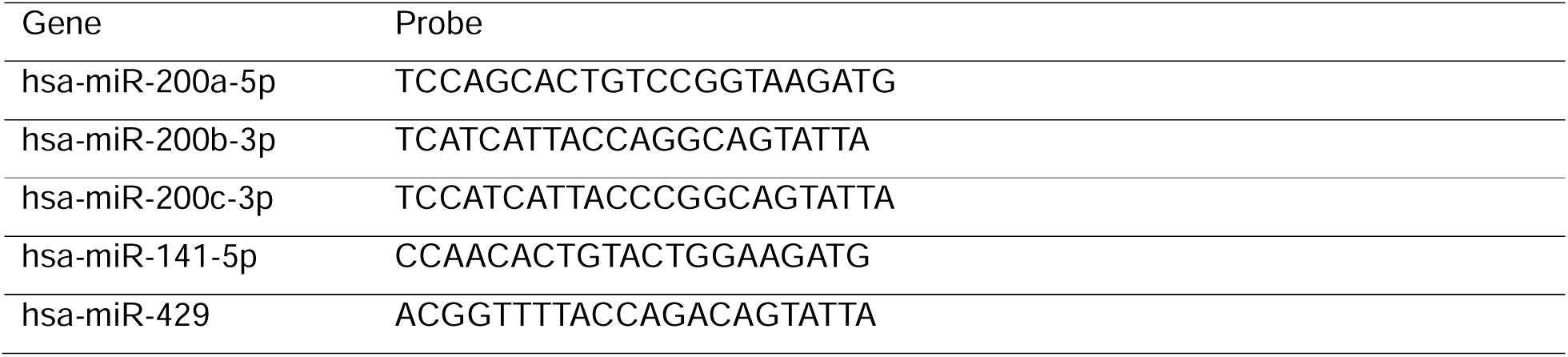
FISH probes.

## 3 RESULTS

### 3.1 Isolation and characterization of the ESC-EVs and FB-EVs

As depicted in Figure 1a, the primary ESCs and fibroblasts (FB) were isolated from human foreskin or fetal rat skin tissues, displaying high proliferative capacity and forming colonies (Fig 1b). Flow cytometry (FCM) analysis revealed the ESCs was positive for integrin CD49f (97.3%) and Krt15 (95.1%), while the FB was positive for Vimentin (98.8%) and negative for Pan-Keratin (99.8%) (Fig 1c).

**FIGURE 1.**
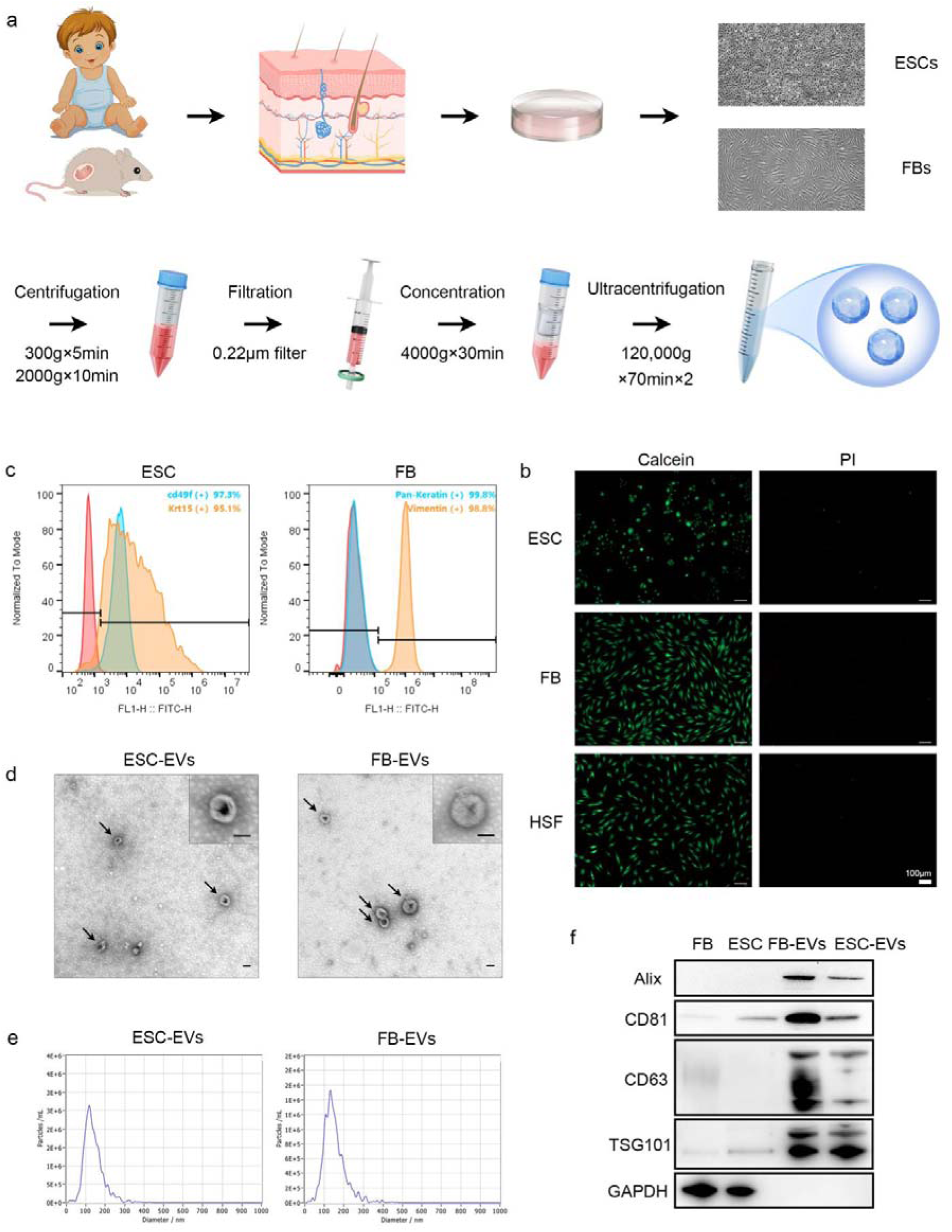
Isolation and characterization of the ESC-EVs and FB-EVs. (a) Schematic diagram of cell separation, culture and the EVs isolation. (b) Calcein and PI stains showed the high proliferative capacity of ESCs, FBs, and HSFs. Scale bar, 100μm. (c) Flow cytometry analysis of ESCs markers (cd49f and Krt15) and FBs markers (Pan-Keratin and Vimentin). (d) Transmission electron microscopy (TEM) images of ESC-EVs and FB-EVs. Scale bar, 500nm (down) and 200nm (up). (e) Nanoparticle tracking analysis of ESC-EVs and FB-EVs from ZetaView. (f) Endosomal (Alix, TSG101) and tetraspanin (CD81, CD63) expression in FBs, ESCs, FB-EVs, and ESC-EVs. GAPDH was used as a negative EVs marker.

Purified EVs were characterized according to morphology, size distribution, surface markers, and surface charge (Fig 1d-f). ESC-EVs and FB-EVs displayed a round shape with a bilayer structure (Fig 1d). Besides, NTA analysis identified that the mean diameters of the ESC-EVs were 127.9nm, and the FB-EVs were 138.7nm (Fig 1e). The protein content in the solution of the ESC-EVs was 0.48μg protein/ml medium, and the FB-EVs was 0.56μg protein/ml medium. The ESC-EVs concentration was 28.4×10^6^ particles/ml medium, which was equivalent to 59.2×10^6^ particles/mg protein, while the FB-EVs was 33.6×10^6^ ml/ medium and identical to 60.0×10^6^ particles/mg protein. In addition, immunoblotting was performed to confirm the presence of typical EV markers Alix, CD9, CD81, CD63, and TSG101 (Fig 1f). These data indicated that the ESC-EVs and FB-EVs we collected were highly purified and suitable for the subsequent experiments.

### 3.2 ESC-EVs were internalized and effectively inducing the MET of HSFs

To determine whether HSFs are recipient cells for ESC-EVs, the ESC-EVs were labeled with PKH67 (green) and the HSF with Actin (red), respectively. After 12 hours of incubation, confocal imaging revealed green fluorescent spots in recipient HSFs, indicating the labeled EVs had been successfully delivered to HSFs (Fig 2a).

**FIGURE 2.**
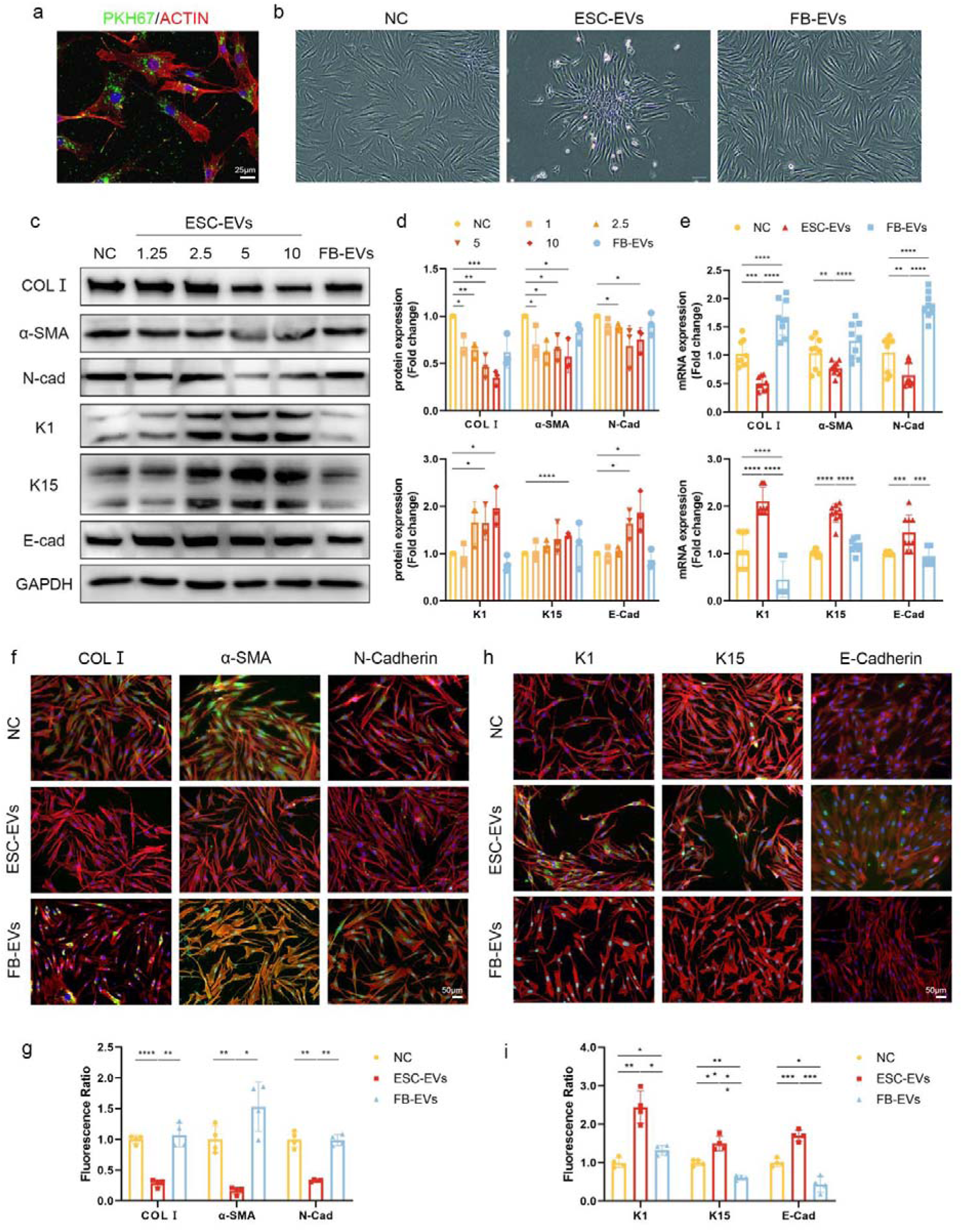
ESC-EVs were internalized and effectively inducing the MET of HSFs. (a) Confocal imaging showed the internalized PKH67-labeled ESC-EVs (green) in ACTIN-labeled HSFs (red). Scale bar, 25μm. (b) Morphologic alterations in HSFs incubated with ESC-EVs, FB-EVs, or PBS for 48h. (c) Western blotting analysis of COLL,α-SMA, N-cad, K1, K15, and E-cad in HSFs incubated with PBS, ESC-EVs at concentrations of 1.25, 2.5, 5, or 10μg/ml or FB-EVs (10μg/ml). (d) Quantification of the relative band density compared to GAPDH. (e) RT-qPCR analysis of the mesenchymal markers COLL, α-SMA, and N-cad, and epithelial markers K1, K15, and E-cad in HSFs treated with ESC-EVs, FB-EVs, or PBS. Graph represented the expression of the markers relative to that of GAPDH. (f) Representative images of COLL,α-SMA, N-cadherin immunofluorescence staining in HSFs incubated with ESC-EVs or PBS for 24h. Scale bar, 50μm. (g) Quantification of the relative fluorescence ratio. (h) Representative images of K1, K15, and E-cadherin immunofluorescence staining in HSFs incubated with ESC-EVs or PBS for 24h. Scale bar, 50μm. (i) Quantification of the relative fluorescence ratio. \**p*<0.05, \*\**p*<0.005, \*\*\**p*<0.0005, \*\*\*\**p*<0.0001.

Then we co-cultured HSFs with different concentrations of ESC-EVs and found that they gradually became irregular and appeared as tufted pavers resembling keratinocytes (Fig 2b). As shown in Fig 2c-d, ESC-EVs significantly decreased the expression of mesenchymal markers in HSFs including collagen I (COLI), α-smooth muscle Actin (α-SMA), and N-cadherin (N-cad), while increasing the expressions of epithelial markers like Krt1 (K1), Krt15 (K15), and E-cadherin (E-cad), and the optimal effective concentration was 5μg/ml (Fig 2d). RT-qPCR analysis and immunofluorescence staining showed the same results (Fig 2e-i). These findings indicated that ESC-EVs effectively induced the MET of HSFs and inhibited its activation to attenuate fibrosis(Ngo *et al*., 2006).

### 3.3 miR-200s were enriched in ESC-EVs and targeting ZEBs in HSFs

To further elucidate the potential mechanisms of ESC-EVs in HSFs’ MET, we detected the miRNA expression profiles between the ESC-EVs and FB-EVs using microarrays (Fig 3a-b). A total of 209 miRNAs were discovered to be significantly different, of which 173 were highly expressed in ESC-EVs and 36 were lowly expressed. Among them, we found that the miR-200 family (miR-200s), including miR-200a, miR-200b, miR-200c, miR-141, and miR-429 were all highly expressed in ESC-EVs, and the differences were significant (LogFC > 1, *p*<0.05, and FDR<0.05). Then we used the clusterProfiler package for Gene ontology (GO) and Kyoto Encyclopedia of Genes and Genomes (KEGG) pathway (Wu *et al*., 2021) annotations, and the ggplot2 package for illustration. As shown in Fig 3c-d, the ZEB1 was enriched in GO analysis, and the ZEB1 and ZEB2 were discovered by KEGG analysis(Huang *et al*., 2019; Bakir *et al*., 2020).

**FIGURE 3.**
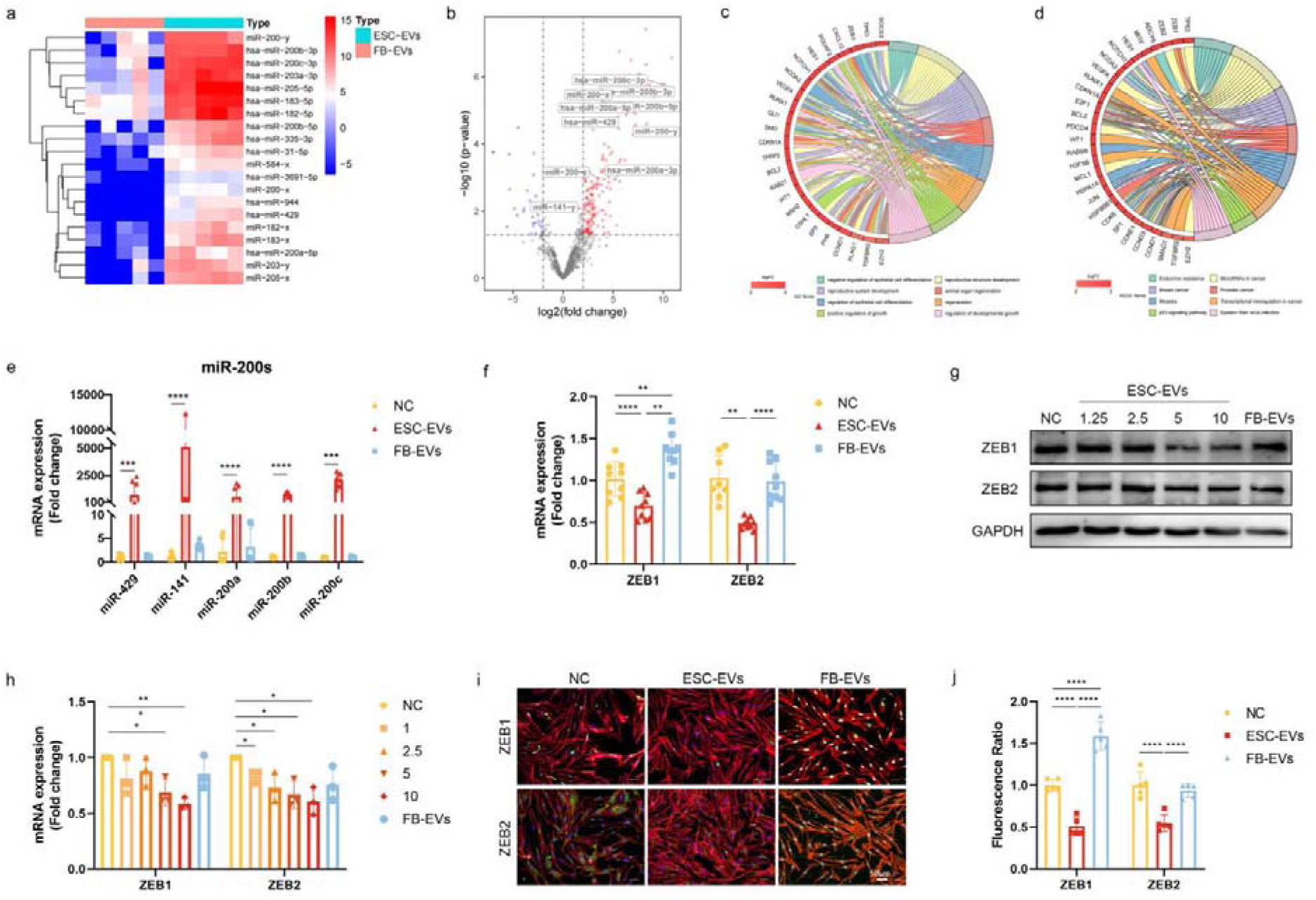
miR-200s were enriched in ESC-EVs and targeting ZEBs in HSFs. (a) Heatmap of the 20 miRNAs with the largest LogFC in the microarray analysis of the miRNA expression profiles between ESC-EVs and FB-EVs. (b) Volcano annotated with miR-200s probes (ESC-EVs vs FB-EVs). Type, High expression (red), Low expression (blue), and No significance (grey). (c) Chord diagram of GO enrichment terms. (d) Chord diagram of KEGG enrichment terms. (e) RT-qPCR analysis of the expression of miR-200s in HSFs treated with ESC-EVs, FB-EVs, or PBS for 24h. Graph represented the expression of the markers relative to that of U6. (f) RT-qPCR analysis of the expression of ZEB1 and ZEB2 in HSFs relative to GAPDH. (g) Western blotting analysis of ZEB1 and ZEB2 in HSFs. (h) Quantification of the relative band density compared to GAPDH. (i) Representative images of ZEB1 and ZEB2 immunofluorescence staining in HSFs. Scale bar, 50μm. (j) Quantification of the relative fluorescence ratio. \**p*<0.05, \*\**p*<0.005, \*\*\**p*<0.0005, \*\*\*\**p*<0.0001.

To investigate whether the miR-200s were successfully transferred to HSFs via ESC-EVs and inhibited ZEBs, we detected the expression of miR-200s in HSFs incubated with PBS, ESC-EVs, and FB-EVs, respectively. After 24 hours of incubation, the expression of miR-200s was remarkably increased in the ESC-EVs group (Fig 3e), while the expression of ZEB1 and ZEB2 was significantly reduced (Fig 3f-j). Taken together, we speculated that ESC-EVs target the inhibition of ZEBs in HSFs by delivering miR-200s.

### 3.4 HS exhibited miR-200s deficiency and ZEBs accumulation

To explicit the expression of miR-200s and ZEBs in human normal skin (NS), hypertrophic scar (HS), and mature scar (MS) tissues, we subsequently examined their expression by fluorescence in situ hybridization (FISH). Compared with the NS, the expression of miR-200s in HS was significantly decreased, while it was restored in MS (Fig 4a-b). At the same time, mIHC also showed the expression of ZEB1 and ZEB2 in the HS was much higher than in NS and MS (Fig 4c-d). These results demonstrated a vital role of the miR-200s/ZEBs axis in the occurrence and development of HS.

**FIGURE 4.**
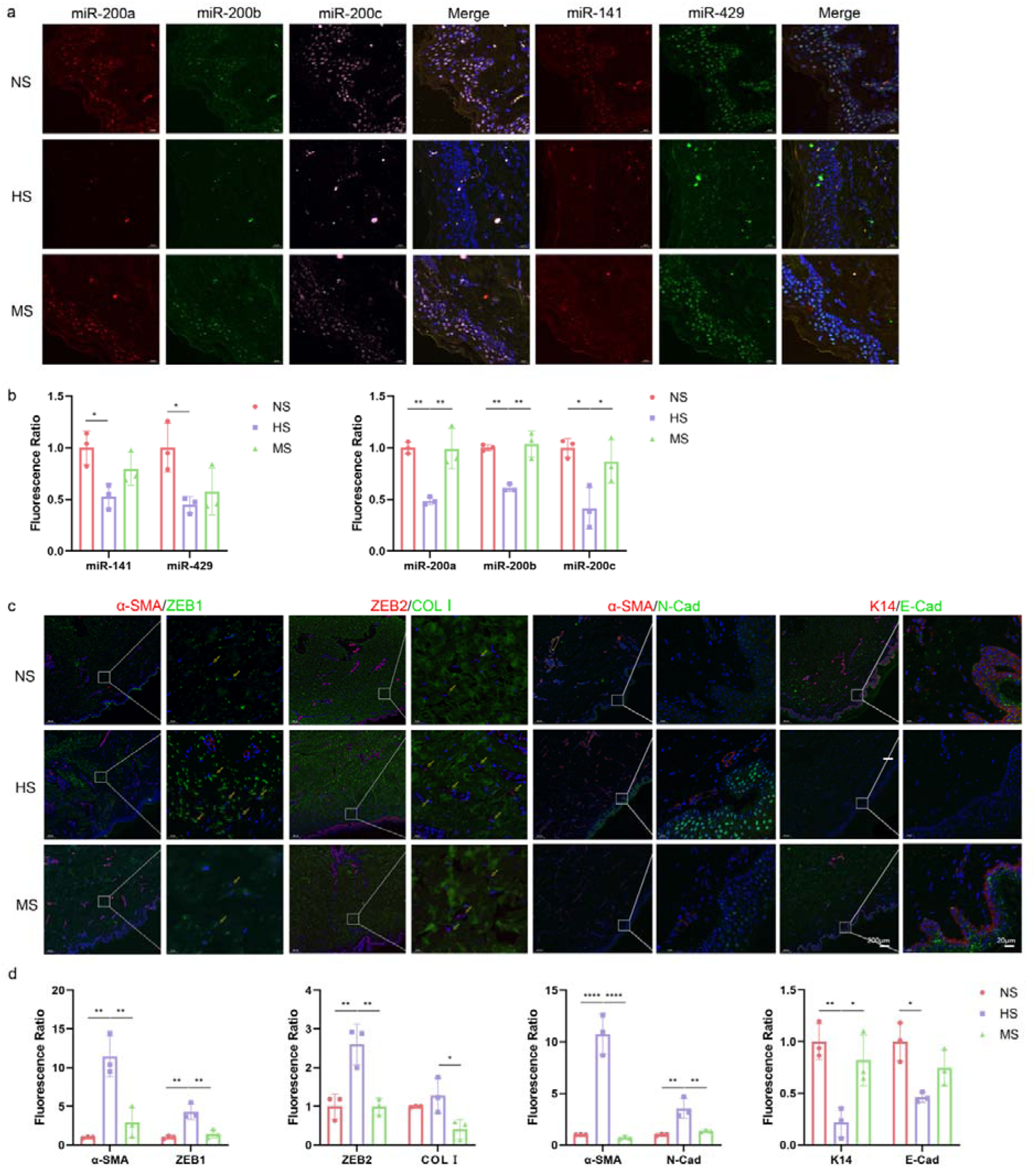
HS exhibited miR-200s deficiency and ZEBs accumulation. (a) FISH stains of miR-200s in normal skin (NS), hyperplasia stage (HS), and mature stage of hypertrophic scar (MS). Scale bar, 20μm. (b) Quantification of the relative fluorescence ratio. (c) mIHC stains of α-SMA, ZEB1, ZEB2, COLL, α-SMA, N-Cad and K14, E-Cad in normal skin, hyperplasia stage, or mature stage of hypertrophic scar. Scale bar, 200μm, 20μm. (d) Quantification of the relative fluorescence ratio. \**p*<0.05, \*\**p*<0.005, \*\*\**p*<0.0005, \*\*\*\**p*<0.0001.

### 3.5 ESC-EVs induced MET of HSFs via miR-200s/ZEBs axis

To further investigate whether the effects of ESC-EVs are dependent on the miR-200s, we targeted all five members of the miR-200s located across two different genomic loci using a multiplexed CRISPR/Cas9 system in a single lentiviral vector based on previous reports(Yu *et al*., 2022). Two DNA fragments encoding guidelines for miR-200b, miR-200a, miR-429 genes on chromosome 1, and miR-200c, miR-141 on chromosome 12, were designed to be deleted in the presence of the four sgRNAs and active Cas9 nuclease (Fig 5a-b). Thus, all miR-200 family members were simultaneously knocked down in ESC-EVs (Fig 5c).

**FIGURE 5.**
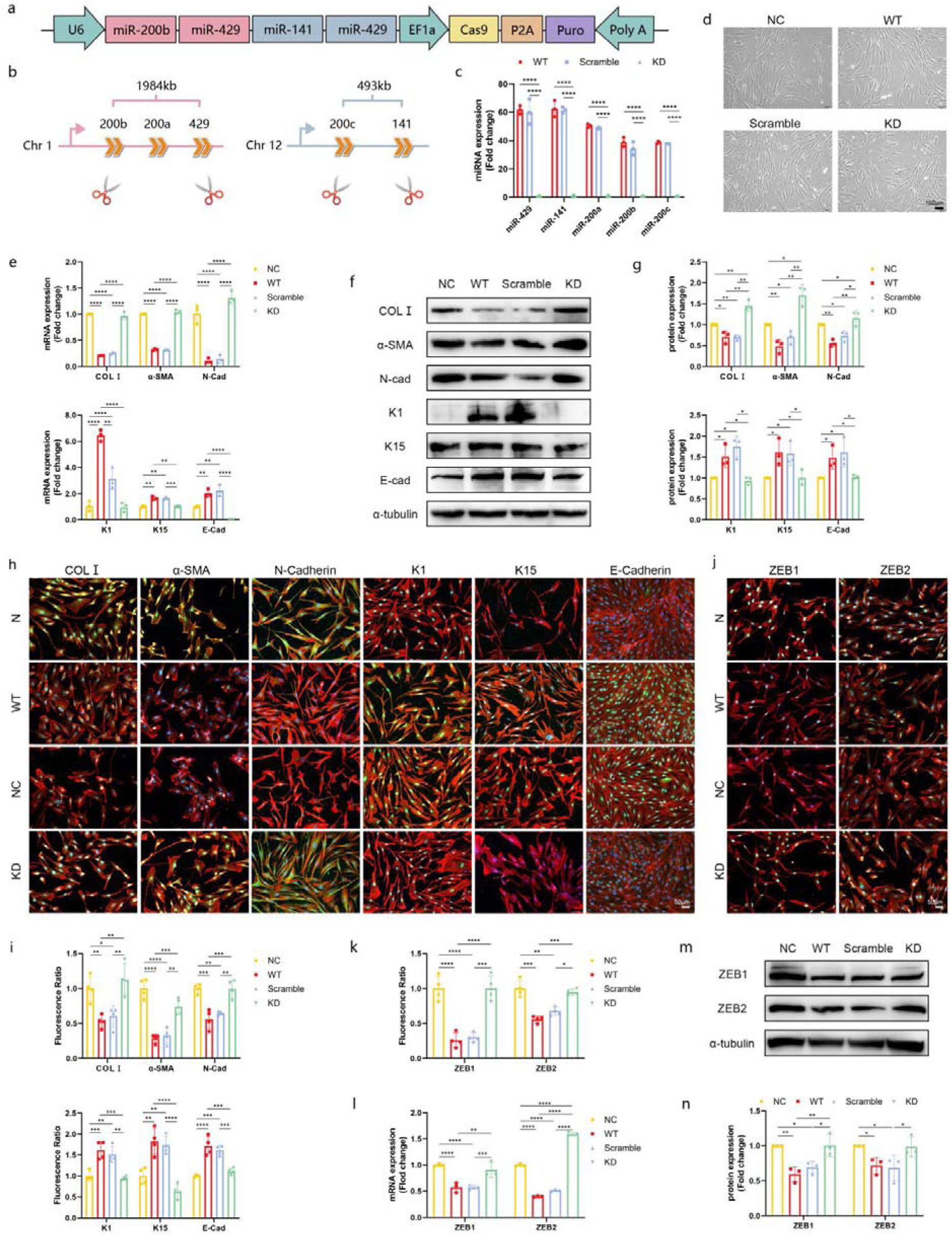
ESC-EVs induced MET of HSFs via miR-200s/ZEBs axis. (a) The multiplex CRISPR/Cas9 in a single lentiviral vector. (b) Five sgRNAs are indicated by scissors targeting two genomic loci of human miR-200 family host genes. (c) RT-qPCR analysis of EVs secreted from wild-type ESC-EVs (WT), ESCs infected with scramble vectors (Scramble), or miR-200s knockdown vectors (KD). (d) Morphologic alterations in HSFs incubated with WT-EVs, Scramble-EVs, KD-EVs, or PBS (NC) for 48h. (e) RT-qPCR analysis of COLL, α-SMA, N-cad, K1, K15, and E-cad in HSFs. Graph represented the expression of the markers relative to that of GAPDH. (f) Western blotting analysis of COLL, α-SMA, N-cad, K1, K15, and E-cad in HSFs. (g) Quantification of the relative band density compared to α-tubulin. (h) Representative images of COLL,α-SMA, N-cad, K1, K15, and E-cad immunofluorescence staining in HSFs incubated with WT-EVs, Scramble-EVs, KD-EVs, or PBS for 24h. Scale bar, 50μm. (i) Quantification of the relative fluorescence ratio. (j) Representative images of ZEB1 and ZEB2 immunofluorescence staining in HSFs. Scale bar, 50μm. (k) Quantification of the relative fluorescence ratio. (l) RT-qPCR analysis of the expression of ZEB1 and ZEB2 in HSFs relative to GAPDH. (m) Western blotting analysis of ZEB1 and ZEB2 in HSFs. (n) Quantification of the relative band density compared to α-tubulin. \**p*<0.05, \*\**p*<0.005, \*\*\**p*<0.0005, \*\*\*\**p*<0.0001.

After being incubated for 24 hours, HSFs cultured with ESC-EVs infected with scramble vectors (Scramble) became tenuous and appeared as tufted pavers, like those incubated with the wild-type ESC-EVs (WT). In contrast, HSFs incubated with ESC-EVs infected with miR-200s knockdown vectors (KD) didn’t show apparent morphological change, aligning with those cultured with the normal medium (N) (Fig 5d). Furthermore, the results from RT-qPCR (Fig 5e), WB (Fig 5f-g), and IF (Fig 5h-i) consistently showed that miR-200s knockdown significantly increased the expression of mesenchymal markers and decreased the expression of epithelial markers. Meanwhile, the knockdown of miR-200s restored the expression of both ZEB1 and ZEB2 suppressed by wild-type ESC-EVs in HSFs (Fig 5j-n). These findings revealed that ESC-EVs induced MET of HSFs via the miR-200s/ZEBs axis.

### 3.6 ESC-EVs-miR-200s suppressed the migration and contractile ability of HSFs

To express the effect of ESC-EVs on HSFs’ function, we detected HSFs’ proliferation, migration, and contractile ability under different treatments. Although CCK8 and Ki67 staining experiments consistently showed that the growth of HSF was not significantly affected by ESC-EVs (Fig 6a-c, Supplementary S1), the scratch test showed that the migration of HSFs was considerably inhibited by ESC-EVs treatments compared to the PBS and FB-EVs groups (Fig 6f). In addition, the collagen gel contraction test also displayed that ESC-EVs significantly inhibited the contractile ability of HSFs while FB-EVs did not.

**FIGURE 6.**
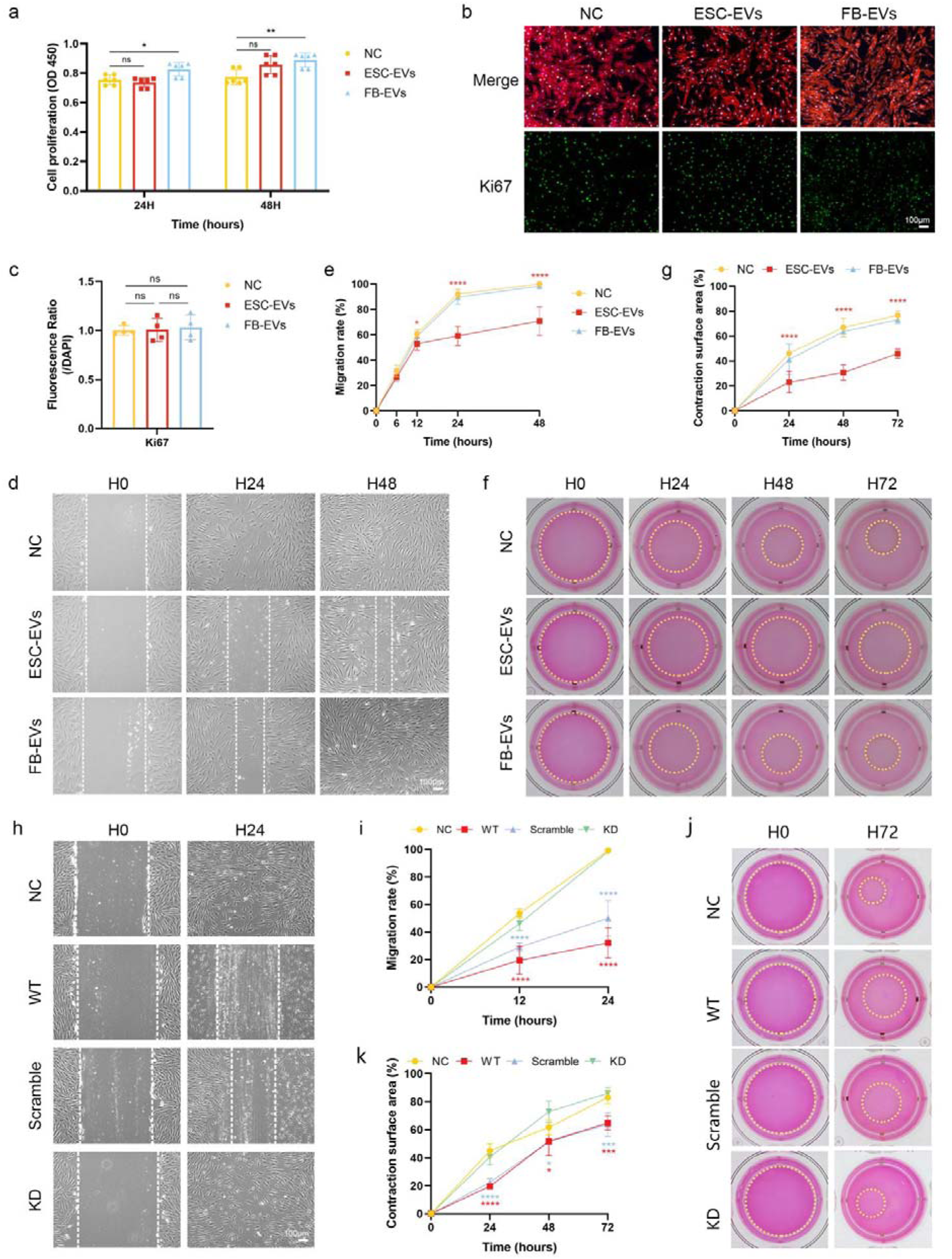
ESC-EVs inhibited the activation of HSFs via miR-200s. (a) CCK8 assay to detect the impact of both ESC-EVs, FB-EVs, or PBS on HSFs cell viability. (b) Representative images of Ki67 immunofluorescence staining in HSFs. Scale bar, 100μm. (c) Quantification of the relative fluorescence ratio. (d) Scratch tests showed the comparison of relative migration areas among ESC-EVs, FB-EVs, or PBS at 0h and 24h. Scale bar, 100μm. (e) Quantification of the migration rate at 6h, 12h, 24h and 48h. (f) Collagen gel contraction test illustrated the contraction of ESC-EVs, FB-EVs, or PBS populated collagen gels at 0h, 24h and 48h. (g) Quantification of contraction surface areas at 24h, 48h, and 72h. (h) Scratch tests showed the comparison of relative migration areas among WT-EVs, Scramble-EVs, KD-EVs, or PBS at 0h and 24h. Scale bar, 100μm. (i) Quantification of the migration rate at 12h and 24h. (j) Collagen gel contraction test illustrated the contraction of WT-EVs, Scramble-EVs, KD-EVs, or PBS populated collagen gels at 0h and 72h. (k) Quantification of contraction surface areas at 24h, 48h, and 72h. \**p*<0.05, \*\**p*<0.005, \*\*\**p*<0.0005, \*\*\*\**p*<0.0001.

However, after the knockdown of the miR-200s in ESC-EVs, both the scratch test and collagen gel contraction test showed that the migration and contractile abilities of HSFs were significantly enhanced compared to those directly treated with ESC-EVs, indicating that the loss of miR-200s in ESC-EVs resulted in the loss of the ability to inhibit HSF activation (Fig 6h-j). These findings revealed that wild-type ESC-EVs induce MET via the miR-200s/ZEBs axis in HSFs.

### 3.7 Application of ESC-EVs alleviated the HS of rat tail (RHS)

To confirm the anti-fibrotic efficacy of ESC-EVs *in vivo,* we established the HS of rat tail (RHS) model according to previously reported (Wang *et al*., 2024). As shown in Figure 7a, a 6×6 mm full-thickness wound was created on the rat tail. Three weeks after the operation, the rat tail defect was replaced by HS.

**FIGURE 7.**
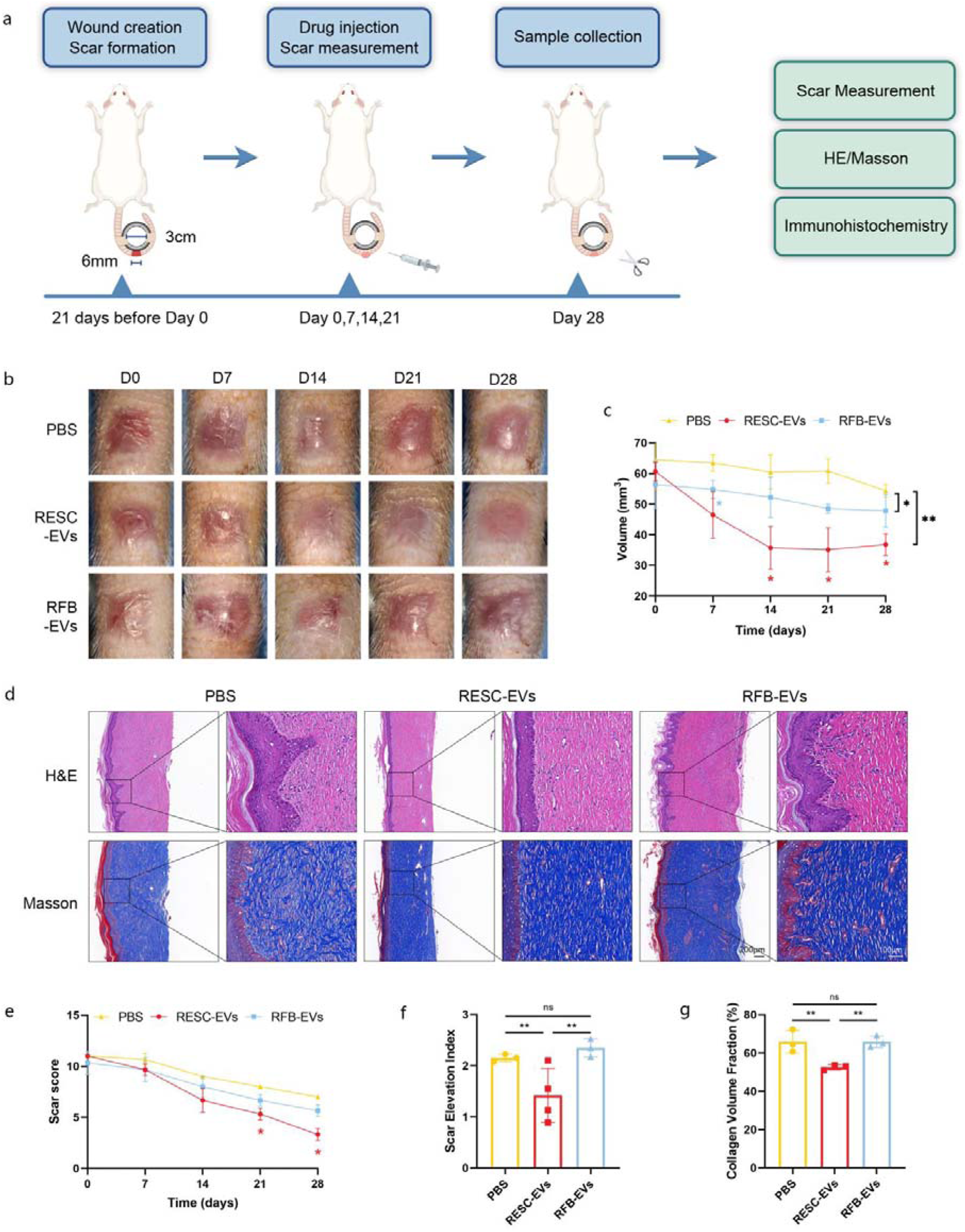
Application of ESC-EVs alleviated the HS of rat tail (RHS). (a) Schematic representation of animal model preparation and administration in RHS. (b) Morphology of RHS on post-injury days 0, 7, 14, 21 and 28. (c) Quantitative evaluation of the scars volume (n = 4 biologically independent samples). (d) Representative images of H&E and Masson’s trichrome stating of RHS sections. Scale bar, 200μm, 100μm. (e) Quantitative evaluation of the scar score (n = 4 biologically independent samples). (f) Quantitative evaluation of the scar elevation index (n = 4 biologically independent samples). (g) Quantitative evaluation of the collagen volume fraction (n = 4 biologically independent samples). \**p*<0.05, \*\**p*<0.005, \*\*\**p*<0.0005, \*\*\*\**p*<0.0001.

ESC-EVs, FB-EVs, or an equal volume of PBS was subcutaneously injected once weekly according to the different groups. No death or abnormality was observed in any animal during the postoperative period. Digital photographs displayed the RHS changes on days 7, 14, 21, and 28 post-wounding (Fig 7b). After two applications of ESC-EVs, the RHS demonstrated a notable decrease in thickness, heightened softness, volume, and a more superficial appearance, thereby contributing to enhanced clinical aesthetics (Fig 7c). Subsequently, H&E and Masson’s trichrome staining revealed a significant reduction in scar elevation index (SEI) and collagen density in the ESC-EVs treated group compared to both control and FB-EVs treated groups, resulting in more organized collagen fibers compared to the FB-EVs and control groups (Fig 7d-g).

### 3.8 ESC-EVs alleviated the RHS through the miR-200s/ZEBs axis

To further investigate the mechanism of ESC-EVs alleviated the RHS, we detected the expression of miR-200s (Fig 8a, b, Supplementary S2a, b), ZEBs, E-Cad, N-Cad, α-SMA, Collagen I, and KRT14 (Fig 8c, d, Supplementary S2c, d). Consistently, the expression levels of α-SMA, N-Cadherin, and Collagen I were significantly reduced, while the miR-200s, Krt14 and E-Cadherin were notably increased in RHS treated with ESC-EVs. In addition, levels of mesenchymal markers exhibited a significant positive correlation, whereas epithelial markers showed a negative correlation with ZEB1/2, consistent with our *in vitro* observations. Taken together, these findings revealed the efficacy of ESC-EVs in attenuating HS by promoting MET of HSFs via the miR-200s/ZEBs.

**FIGURE 8.**
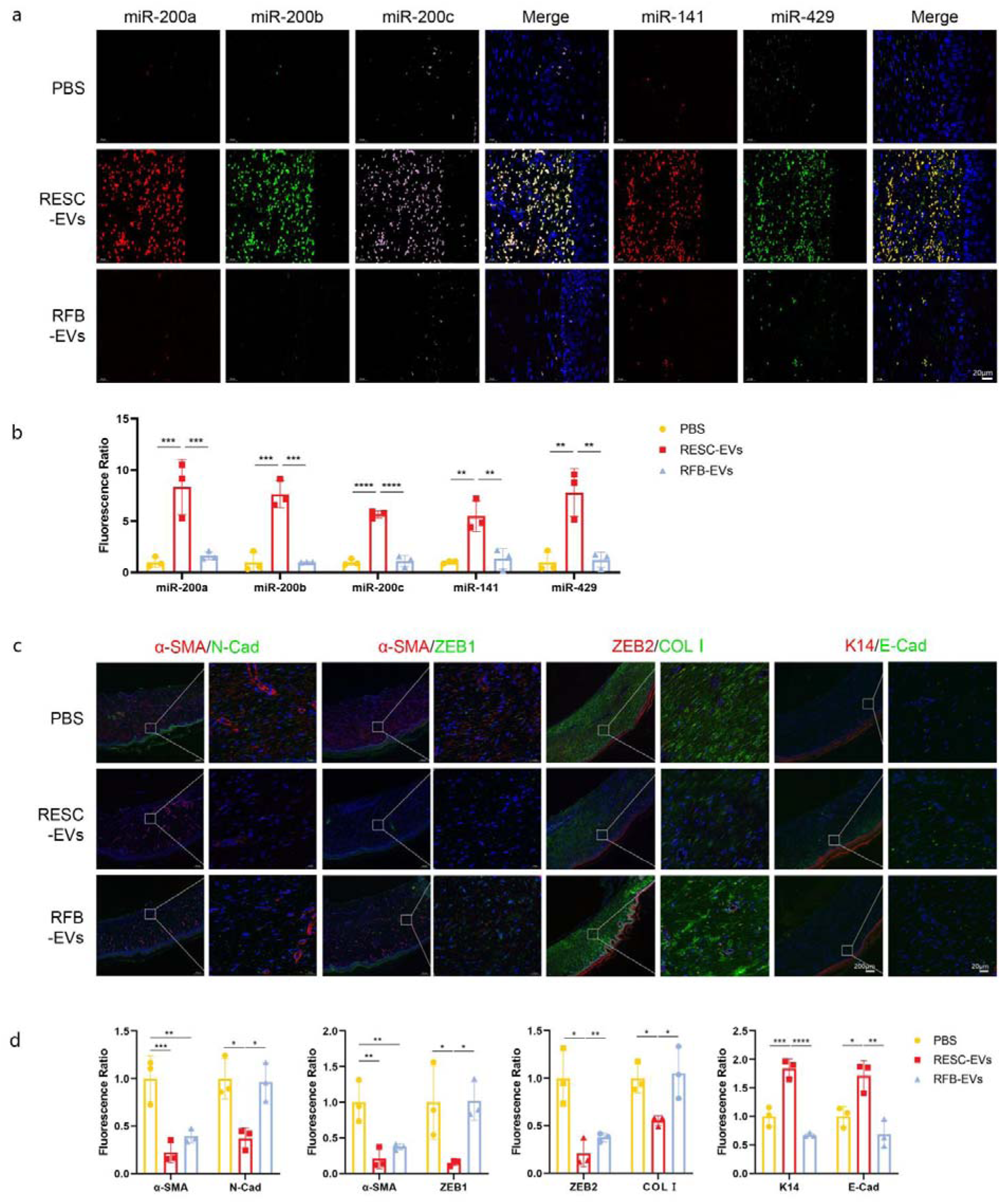
ESC-EVs alleviated the RHS through the miR-200s/ZEBs axis. (a) FISH stains of miR-200s in PBS, RESC-EVs, RFB-EVs treated RHS. Scale bar, 20μm. (b) Quantification of the relative fluorescence ratio. (c) Representative images of COLL,α-SMA, N-cad, K1, K15, E-cad, ZEB1 and ZEB2 immunofluorescence staining. Scale bar, 200μm, 20μm. (d) Quantification of the relative fluorescence ratio. \**p*<0.05, \*\**p*<0.005, \*\*\**p*<0.0005, \*\*\*\**p*<0.0001.

**FIGURE 9.**
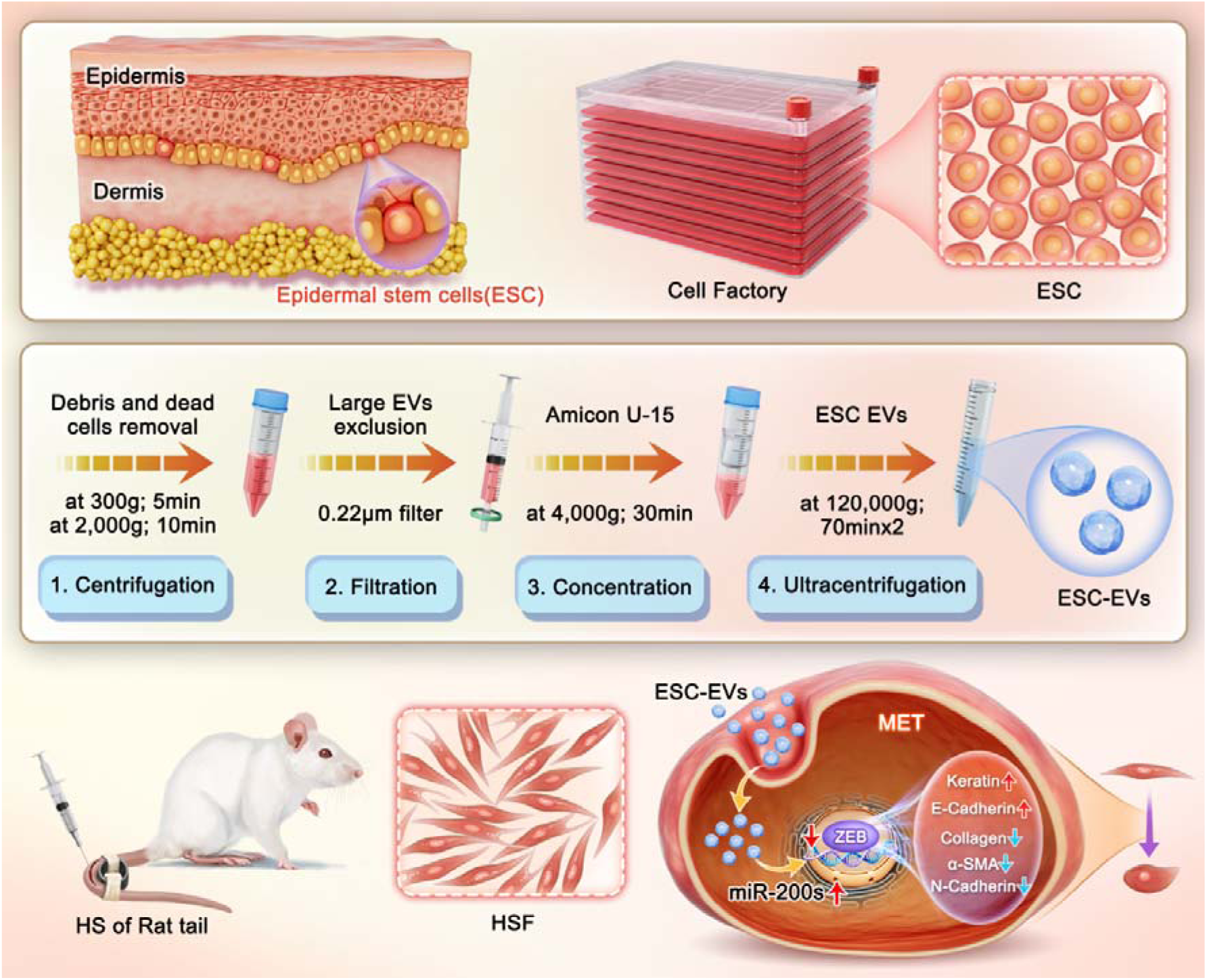
Schematic illustration of ESC-EVs induced HSFs’ MET to alleviate HS via the miR-200s/ZEBs axis. Epidermal stem cells (ESC) were cultured in cell factories to obtain sufficient extracellular vesicles (ESC-EVs). EVs were isolated using a differential ultracentrifugation method. Both in vitro cell experiments and in vivo rat tail HS (RHS) models thoroughly validated that ESC-EVs were internalized and effectively inducing the MET of HSFs and inhibited its activation to attenuate fibrosis via the miR- 200s/ZEBs axis.

## 4 DISCUSSION

The overproduction of hypertrophic scar fibroblasts (HSFs) caused by excessive EMT is one of the most critical factors leading to HS (Yuan *et al*., 2019). Effectively inhibiting EMT and even inducing MET is significant for treating HS. In the present study, we first found that ESC-EVs could effectively induce the MET of HSFs and inhibit their biological activity. Then, we further elucidated that this therapeutic effect is mediated by the miR-200s encapsulated in ESC-EVs, which targeted and regulated ZEB1 and ZEB2 in HSFs. This vital role and mechanism have been fully validated in both in vitro cell experiments and in vivo RHS models. These findings highlight the anti-fibrosis potential of ESC-EVs and their role in treating skin fibrosis disease.

As a formidable fibrotic skin condition, HS poses an enduring challenge in clinical practice worldwide (Finnerty *et al*., 2016; R *et al*., 2021). Although traditional treatment methods have progressed in recent years, most current first-line treatments for HS are still unsatisfactory with many side effects (Ogawa *et al*., 2021). To develop novel, practical therapeutic approaches explicitly focusing on its pathogenesis is imperative.

ESCs represent a pluripotent stem cell population capable of proliferating into specialized keratinocytes and skin appendages (Blanpain and Fuchs, 2006). Residing in the basal layer of the epidermis, ESCs interface intimately with the underlying basement membrane, thereby encountering a milieu of dermal cells (Fujiwara *et al*., 2018). Its biological behavior plays a vital role in the differentiation of fibroblasts during wound healing and scar formation (Wang *et al*., 2017b). Our previous studies have validated its capacity to enhance impaired wound healing and relieve HS (Huang *et al*., 2020). However, as cellular therapy remains beset by challenges, including cellular immunogenicity, storage stability, ethical considerations, and risks associated with embolism and tumorigenesis, the clinical research and application of ESCs have been greatly limited (Herberts *et al*., 2011).

As central cell-cell communication mediators, EVs share similar functions with the maternal cells (Théry *et al*., 2018). Compared with stem cell transplantation, EVs therapy has the advantages of low immunogenicity, high safety, easy storage and management, and mass production, which is an ideal choice for clinical application (Lener *et al*., 2015). Recent studies have spotlighted the promising potential of EVs derived stem cells, such as mesenchymal stem cells (Helissey *et al*., 2022; Zhao *et al*., 2022) and adipose stem cells (Yang *et al*., 2024; Zhou *et al*., 2024) in mitigating and treating fibrotic diseases (Yang *et al*., 2022). Nevertheless, as the most critical progenitor cell for skin regeneration, the vital role of EVs derived from the primary cultured ESCs has not been thoroughly investigated (Duan *et al*., 2020).

Herein, we established an efficient ESC-EVs isolation and application system and found it significantly inhibited the biological activity of HSFs in vitro and in vivo. ESC-EVs not only inhibited HSFs’ migration and contractility, but also prompted a noteworthy MET phenotypic shift in HSFs, marked by suppressed COLL, α-SMA, and N-cadherin expression, alongside enhanced K1, K15, and E-cadherin expression. On the RHS models, ESC-EVs treated groups consistently displayed significant MET tendency and fibrosis alleviation. Therefore, ESC-EVs exhibited great potential in catalyzing MET, curtailing HSF activation, and ameliorating abnormal fibrosis within HS.

To investigate the mechanism of ESC-EVs on HS suppression, we further sequenced ESC-EVs and found 173 miRNAs highly expressed miRNAs compared to FB-EVs. Notably, members of the miR-200 family (miR-200s)—miR-200a, miR-200b, miR-200c, miR-141, and miR-429—ranked prominently among the top 20 differentially expressed miRNAs. These results align with previous studies that prompt the vital involvement of miR-200s in attenuating fibrotic processes (Liu *et al*., 2012; Li *et al*., 2014). Then, we assessed the miR-200s expression in normal human skin, HS, and mature scars by fluorescence in situ hybridization (FISH) and found that miR-200s levels in HS were significantly decreased compared to normal skin, while a deficit resolved during scar maturation. However, When the ESC-EVs were applied to the HSF or RHS, notably elevated miR-200s could be observed, which reminded us that ESC-EVs-miR-200s play a vital role between the crosstalk of ESCs and HSFs.

To clarify the target of ESC-EVs in HSFs, GO/KEGG analysis of upregulated genes showed highlighted enrichment in pathways involving ZEB1 and ZEB2, paralogs belonging to the zinc-finger E-box-binding homeobox (ZEB) transcription factor family known for promoting EMT (Bakir *et al*., 2020). Our investigation consistently confirmed heightened obvious ZEB1 and ZEB2 expression in HS compared to normal skin and mature scars. Since extensive literature supports miR-200s’ role in suppressing EMT by targeting ZEB1/2 (Burk *et al*., 2008; Diaz-Riascos *et al*., 2019; Bhome *et al*., 2022), we highly speculated that ESC-EVs targeted ZEBs through miR-200s. Notably, applying ESC-EVs effectively elevated miR-200s and mitigated ZEB1 and ZEB2 expression in HSFs, ultimately inducing its MET and alleviating HS. However, when we knocked down all five members of the miR-200s using a multiplexed CRISPR/Cas9 system, the effect of ESC-EVs on HSFs was significantly weakened, demonstrating the effects of ESC-EVs on HSFs’ MET are mainly dependent on the miR-200s/ZEBs axis.

This study has its limitations. It primarily focused on EVs’ small RNA cargo, overlooking other potentially influential components. In addition, although the HSFs are the most critical targeted cell, other inducements, such as angiogenesis and immune regulation, should be explored in the future. However, the primary objective of this project was to provide evidence of novel ESC-EVs that relieve HS, explore its most critical mechanisms, and identify avenues for future investigation. To our knowledge, this study innovatively revealed that ESC-EVs can induce MET, attenuate HSF activation, and mitigate fibrosis in vitro and in vivo by modulating ZEB1 and ZEB2 through miR-200s. These findings provided a novel therapeutic strategy and elucidated the mechanism for using ESC-EVs and miR-200s as a clinical treatment of HS and other fibrotic disorders.

## 5 CONCLUSION

In summary, our study highlights a significant breakthrough in understanding the role of ESC-EVs in alleviating HS and uncovers a previously unknown mechanism by which ESC-EVs could induce HSFs’ MET via miR-200s/ZEBs axis. As soon as uptaken by HSF, the ESC-EVs-miR-200s directly targeted to ZEB1/2 and induced the MET of HSF, ultimately attenuating HSFs’ biological functions and alleviating HS. Our findings are expected to provide novel targets and strategies for the precise clinical treatment of HS and other skin fibrosis diseases.

## ACKNOWLEDGEMENTS

This work was mainly supported by the National Natural Science Foundation of China (No. 82002043, 82172207, 82273561), Natural Science Foundation of Guangdong Province (No. 2023A1515010265, 2023A1515010146, 2021A1515011806), Guangzhou Municipal Science and Technology Project (No. 2025A04J4078).

## DATA AVAILABILITY STATEMENT

All data are available in the main text and/or the Supplementary materials, tables and figures. Datasets related to this article can be found in the National Center for Biotechnology Information Gene Expression Omnibus database, under the study accession number GSE287895.

