## Supplemental Figure for "Epidermal stem cell-derived extracellular vesicles induce fibroblasts mesenchymal-epidermal transition to alleviate hypertrophic scar via the miR-200s/ZEBs axis"

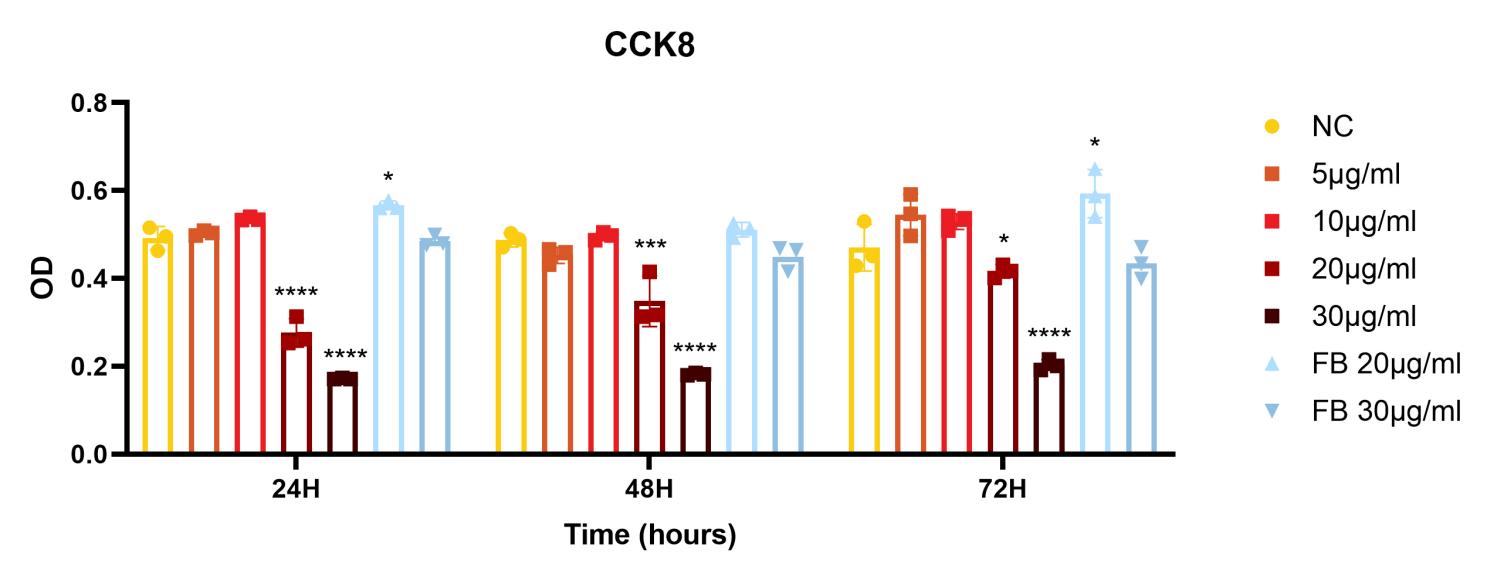

**Supplementary S1** The proliferation of different concentrations of ESC-EVs or FB-EVs treated HSFs was analyzed using CCK-8 assay. **p*<0.05, ****p*<0.0005, *****p*<0.0001.

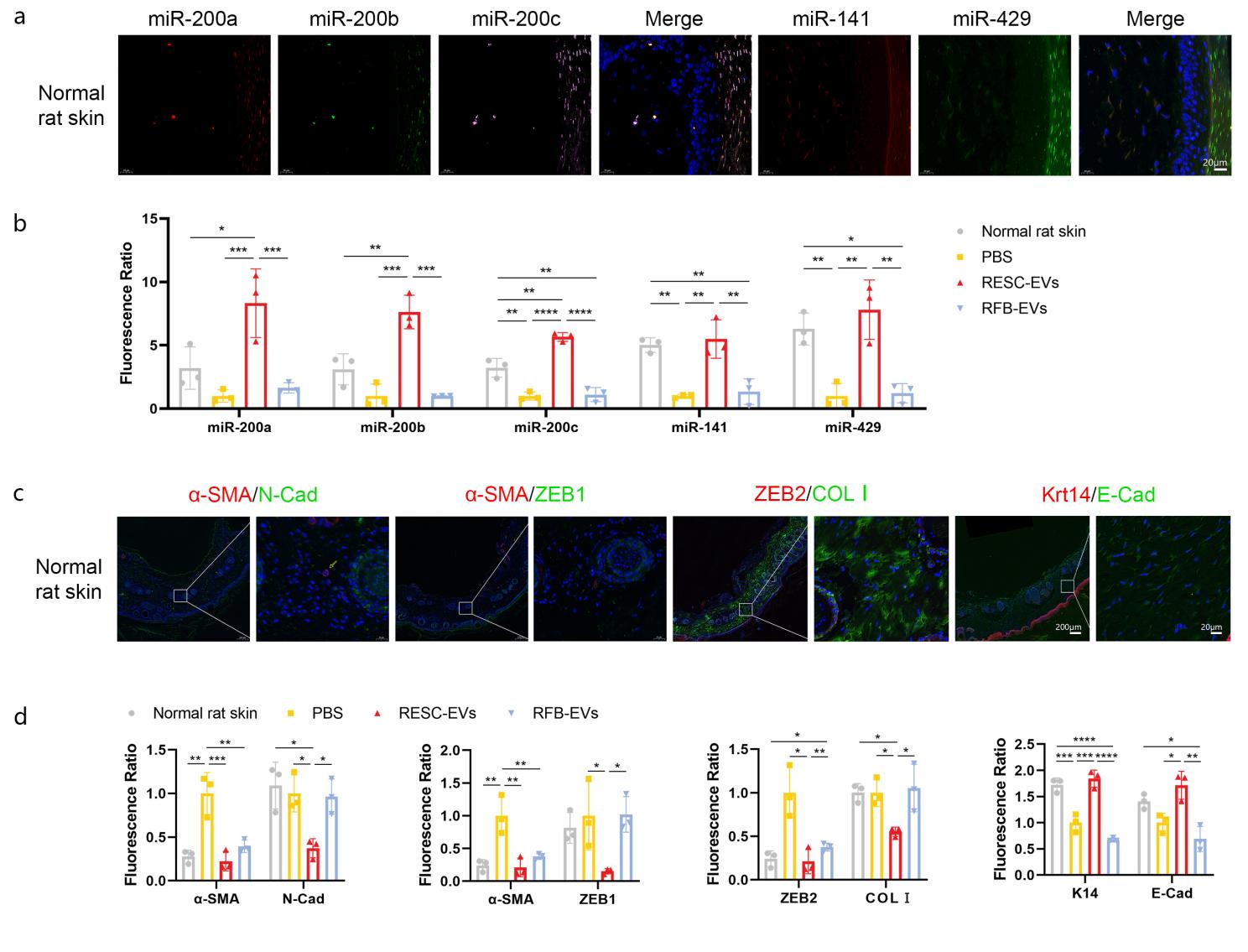

**Supplementary S2** The expression of miR-200s/ZEB axis and MET related indicators in normal rat tail skin. (a) FISH stains of miR-200s in normal rat skin. Scale bar, 20μm. (b) Quantification of the relative fluorescence ratio. (c) Representative images of COLⅠ,α-SMA, N-cad, K1, K15, E-cad, ZEB1 and ZEB2 immunofluorescence staining. Scale bar, 200μm, 20μm. (d) Quantification of the relative fluorescence ratio. **p*<0.05, ***p*<0.005, ****p*<0.0005, *****p*<0.0001.
